# STAT3 operates as an inflammation-dependent transcriptional switch

**DOI:** 10.1101/2025.09.26.678857

**Authors:** Simon Grassmann, Hyunu Kim, Christin Friedrich, Marine Pujol, Sherry Fan, Jennifer Zhang, Giorgi Beroshvili, Veit R. Buchholz, Georg Gasteiger, Joseph C. Sun

**Affiliations:** Immunology Program, Memorial Sloan Kettering Cancer Center, New York, NY 10065, USA; Immunology and Microbial Pathogenesis Program, Weill Cornell Medical College, New York, NY, 10065, USA; Louis V. Gerstner Jr. Graduate School of Biomedical Sciences, Memorial Sloan Kettering Cancer Center, New York, NY 10065, USA; Wuerzburg Institute of Systems Immunology (WueSI), Max Planck Research Group at the Julius-Maximilians-Universitaet Wuerzburg, Germany; Division of Immunology, German Cancer Research Center (DKFZ), Heidelberg, Germany; Institute for Medical Microbiology, Immunology and Hygiene, Technische Universitaet Muenchen (TUM), Munich, Germany

## Abstract

Signal transducer and activator of transcription 3 (STAT3) is a key regulator of immune cell function, but its role in lymphocytes remains incompletely understood. Here, we show that STAT3 has a context-dependent function in antiviral natural killer (NK) cells, either promoting or impairing adaptive NK cell responses dependent on the level of inflammation. STAT3 is recruited to distinct genomic sites under homeostatic versus inflammatory environments, where it drives different transcriptional programs. Through this re-localization, STAT3 regulates downstream transcription factors MYB and BLIMP-1 in an inflammation-dependent manner to shape NK cell differentiation under homeostasis and during infection. Thus, STAT3 acts as a transcriptional switch that integrates cytokine signals to control lymphocyte adaptation to different environments. This mechanism highlights how therapeutic interventions targeting STAT3 can result in different outcomes depending on the degree of inflammation.

**HIGHLIGHTS:**

- STAT3 exerts a context-dependent role on adaptive NK cells during viral infection
- STAT3 modulates IL-15 signaling by competing with STAT5 and regulating MYB
- Homeostatic versus inflammatory cytokines relocate STAT3 to distinct genomic sites
- Downstream transcription factors MYB and BLIMP-1 govern STAT3-dependent differentiation

## INTRODUCTION

Modulating immune cell function is a promising therapeutic strategy in autoimmunity, infection, and cancer. Immune function is closely tied to cell differentiation, which is largely shaped by receptor–ligand interactions and cytokine signals. Therapeutic interventions often exploit these pathways: immune checkpoint blockade prevents adoption of an “exhausted” state by targeting PD-1 and CTLA-4, while cytokine pathways are blocked to restrain autoreactive cells in autoimmunity^1,2^. Despite success in autoimmunity, targeting cytokines in infectious disease and cancer has yielded only moderate benefit^3^. A central challenge is the redundancy and context-dependent nature of many cytokines and their downstream transcription factors^4^; for example, diverse inflammatory, homeostatic, and immunosuppressive cytokines converge on the same JAK/STAT pathways. Dissecting the mechanistic basis of this context dependence is critical for developing effective cytokine-targeted therapies.

Cytotoxic lymphocytes comprise CD8^+^ T cells and NK cells and are critical for host defense against virus infections and cancer. Cytokines regulate the balance of effector versus adaptive differentiation in cytotoxic lymphocytes. Inflammatory cytokines such as type I interferon (IFN) and interleukin (IL)-12 activate STAT1/STAT2 and STAT4, respectively, to promote effector differentiation^5–8^. In contrast, STAT5 is activated by cytokines of the common gamma-chain family, is crucial for homeostatic maintenance, and promotes proliferation while maintaining stemness in lymphocytes^9–11^. Compared to these STATs, STAT3 plays a more complex and less defined role^12^. Traditionally viewed as downstream of immunosuppressive IL-10, STAT3 is thought to inhibit or suppress lymphocyte functions^13,14^. However, there are conflicting reports indicating that STAT3 can also promote lymphocyte proliferation and differentiation^15,16^. Thus, STAT3 represents the canonical example of a context-dependent transcription factor downstream of cytokines in cytotoxic lymphocytes.

Here, we investigate the context-dependent role of STAT3 in NK cells using the mouse cytomegalovirus (MCMV) infection model, where NK cells display adaptive features including antigen specificity, clonal expansion, and memory formation^17^. Prior work demonstrated that STAT1^18^, STAT2^19^, STAT4^20^, and STAT5^10^ are required to promote adaptive NK cell responses in MCMV infection. In contrast, here we find that STAT3 can both inhibit or promote adaptive NK cell responses depending on viral dose. Mechanistically, STAT3 genomic binding is redirected by homeostatic versus inflammatory cytokines, engaging distinct transcriptional networks—particularly MYB and BLIMP-1. Our findings provide a molecular framework for cytokine context-dependence and identify STAT3 as a key mediator of this phenomenon in innate lymphocytes.

## RESULTS

### STAT3 exerts a context-dependent role in adaptive responses of antiviral NK cells

In response to MCMV infection, Ly49H^+^ NK cells mount adaptive responses^17,21,22^ However, the level of NK cell expansion does not scale with virus inoculum dose. Instead, we observed that infection with a high dose of MCMV (5,000 PFU) leads to less expansion of Ly49H^+^ NK cells than low-dose infection (1,000 PFU) (**Fig. 1A** and **Supp. Fig. 1A-B**). To understand why NK cells expand less in high- versus low-dose infection, we performed RNA sequencing (RNAseq) from WT NK cells on day 2 post-infection (PI). Among the top differential pathways in GSEA analysis we found a pathway representing STAT3 targets (**Fig. 1B**). Thus, we hypothesized that STAT3 might regulate NK cell expansion in low- versus high-dose infection. To directly test this hypothesis, we co-transferred equal numbers of NK cells from *Ncr1^Cre^* x *Stat3^flox/flox^*mice (*NK-Stat3^-/-^,* CD45.2) and WT mice (CD45.1) into *Ly49h^-/-^* hosts subsequently infected with MCMV. STAT3-deficient NK cells showed a defect in adaptive expansion during low-dose infection with MCMV compared to WT controls (**Fig. 1C**, left). However, in high-dose infection, the role of STAT3 in antiviral NK cells was reversed, as STAT3-deficient NK cells showed better expansion than WT counterparts (**Fig. 1C**, right). To further characterize WT versus STAT3-deficient NK cells in low- versus high-dose infection, we performed transcriptomic analysis at different time points following infection (**Fig. 1D** and **Supp. Fig. 1C-E**). Interestingly, we found a mostly negative correlation in the expression pattern of genes differentially regulated between WT and *NK-Stat3^-/-^* NK cells in low- versus high-dose infection (**Fig. 1E**).

**Figure 1:**
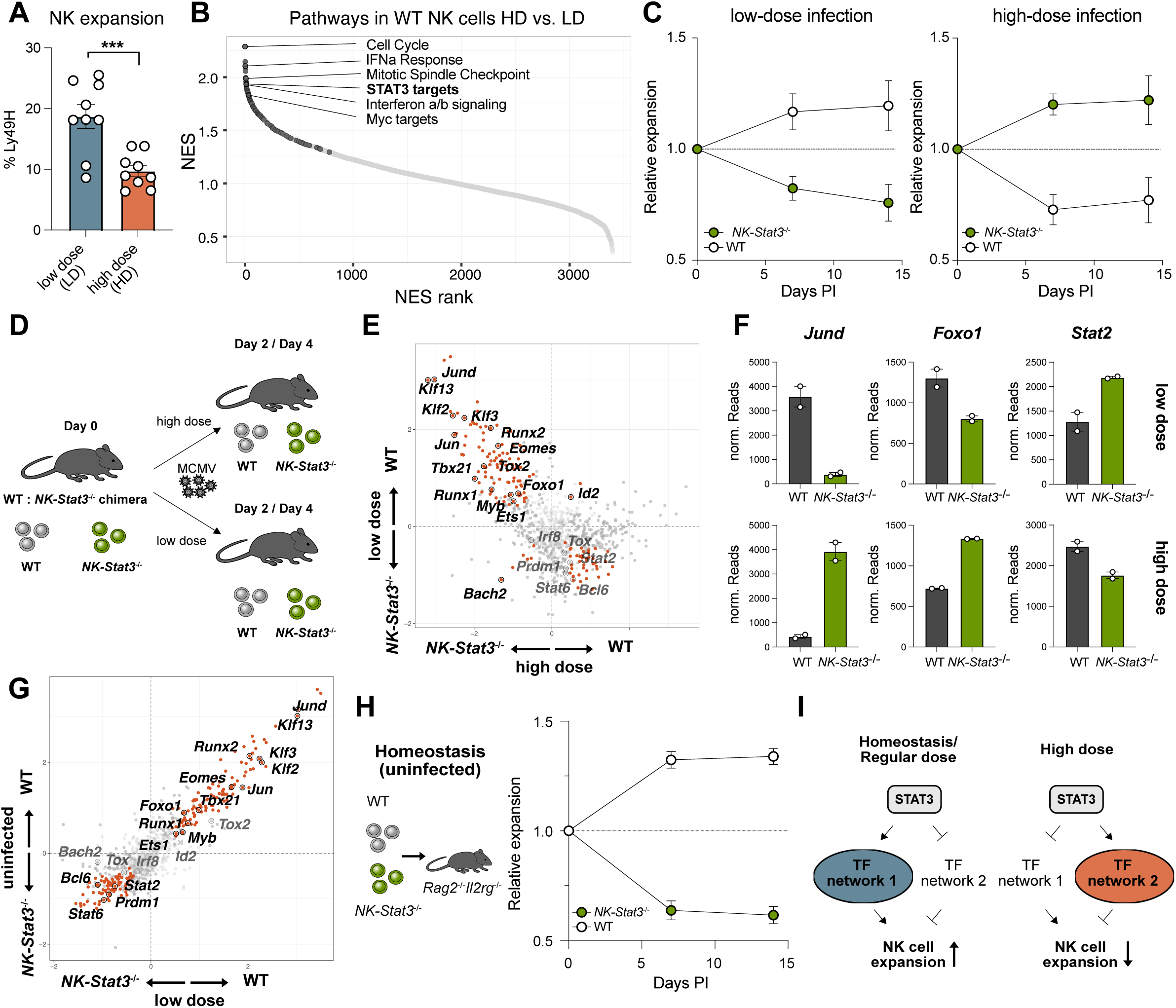
STAT3 exerts a context-dependent role in antiviral NK cells during low- and high-dose MCMV infection. **(A)** Transfer of Ly49H^+^ NK cells into *Ly49h^-/-^*mice and measurement of expansion in spleens on day 7 following infection with a low dose (LD) versus high dose (HD) of MCMV. **(B)** Normalized Enrichment Score (NES) from GSEA analysis of differential gene expression contrasting HD versus LD infection on day 2 PI of WT NK cells, subsetted on positively enriched (NES > 0) pathways. Circles in color show statistically significant enriched pathways (padj ≤ 0.05). **(C)** Co-transfer of WT and *NK-Stat3^-/-^*NK cells into *Ly49h^-/-^* mice infected with a LD (left graph) or HD (right graph) of MCMV and consecutive analysis in blood. **(D)** Schematic of RNA-seq experiment in competitive WT:*NK-Stat3^-/-^* chimera infected with LD or HD. **(E)** Scatterplot of WT versus *NK-Stat3^-/-^*NK cell transcriptomic changes (log2 fold change) in HD (x-axis) and LD (y-axis) of MCMV infection. Circles in color represent significant differentially expressed (DE) genes in both dose conditions. **(F)** Normalized Reads for selected transcription factors shown in E). **(G)** Scatterplot of WT versus *NK-Stat3^-/-^* NK cell transcriptomic changes (log2 fold change) in LD group (x-axis) versus uninfected controls (y-axis). **(H)** Co-transfer of WT versus *NK-Stat3^-/-^*NK cells into *Rag2^-/-^ x Il2rg^-/-^* mice and measurement of expansion in blood. **I)** Model shows instruction of distinct transcription factor networks by STAT3 in uninfected, low-dose, and high-dose MCMV infection. Data in 1A) and 1C are pooled from 2-3 experiments. Statistics in 1A is derived from unpaired t-test.

STATs are transcription factors (TFs) that instruct downstream TF networks to guide immune cell differentiation^23^. To assess if STAT3 engages distinct transcription factor networks in low- and high-dose infection, we compared expression of TFs between WT and *NK-Stat3^-/-^* NK cells during infection (**Fig. 1E-F**). Indeed, many TFs showed an opposite role for STAT3 in low- versus high-dose infection (**Fig. 1E-F**). Among the TFs upregulated by STAT3 in low-but not high-dose infection, we found ones crucial for adaptive lymphocyte programming such as *Klf2*, *Foxo1* and *Myb*.

We observed that STAT3-deficient NK cells are already distinct from WT NK cells under steady state (**Supp. Fig. 1F**). Comparing the TF networks engaged by STAT3 in different contexts revealed that regulation of similar transcription factors was positively correlated under steady state and during low-dose infection (**Fig. 1G** and **Supp. Fig. 1G**). This difference was not due to an insufficient infection, as infection-dependent transcriptomic changes were larger than differences between WT and *NK-Stat3*^-/-^ NK cells (**Supp. Fig.1C**). Accordingly, *NK-Stat3^-/-^* NK cells showed a homeostatic proliferation disadvantage reminiscent of our observation in low-dose infection even in uninfected *Rag2^-/-^ x Il2rg^-/-^* mice (**Fig. 1H**). Thus, STAT3 appears to exert a context-dependent role in NK cells, coinciding with the engagement of distinct TF networks (**Fig. 1I**), where during homeostasis and low-dose infection, STAT3 regulates a transcription factor network that enhances NK cell expansion. However, during high-dose infection, STAT3 regulates expression of distinct transcription factors linked to impaired NK cell adaptive responses.

### STAT3 modulates IL-15 signaling during homeostasis to promote proliferation

A previous study reported that STAT3-deficient NK cells had no pronounced phenotypic changes at steady state using flow cytometry^13^, an observation we independently confirmed (**Supp. Fig. 2A-C**). However, *NK-Stat3^-/-^* NK cells analyzed by RNA-seq were distinct from WT NK cells (**Fig. 2A**). Genes differentially expressed in WT versus *NK-Stat3^-/-^* NK cells were enriched for crucial pathways such as MYC and MYB targets, IL-2 signaling, and cell cycle (**Fig. 2B**). Homeostatic NK cell maintenance is predominantly regulated by IL-15^24^. Although previous reports have suggested that many cytokines, including IL-15 and type-I interferon, can result in STAT3 phosphorylation in other cell types^25–28^, we only observed robust pSTAT3 after stimulation of primary NK cells with IL-10 (**Supp. Fig. 2D-F**). To test whether STAT3 activation alters IL-15 dependent NK cell proliferation, we stimulated NK cells with IL-10 and IL-15 and measured proliferation via CTV dilution. Whereas IL-10 did not affect proliferation at high IL-15 concentrations, at lower (and likely more physiologic) IL-15 levels IL-10 profoundly enhanced proliferation in WT but not *NK-Stat3^-/-^* NK cells (**Fig. 2C**-**D**). Thus, our data suggests that the positive role of STAT3 for NK cell proliferation under homeostasis and during low dose infection was likely due to an interaction of STAT3 with IL-15 signaling.

**Figure 2:**
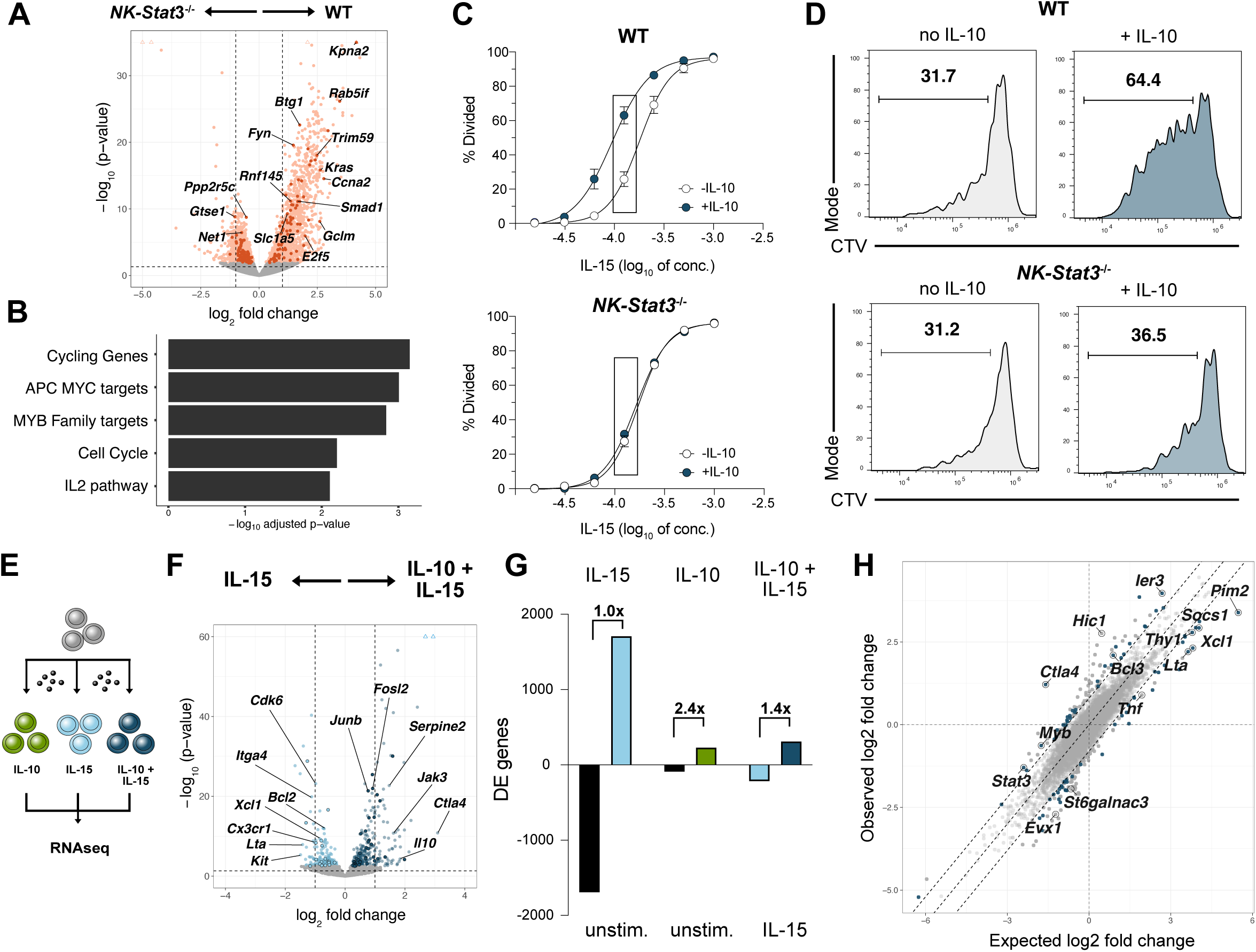
STAT3 activation modulates IL-15 signaling to drive proliferation of NK cells under homeostasis. **(A)** Volcano plot of gene expression for WT versus *NK-Stat3^-/-^*NK cells in uninfected mice (from competitive WT:*NK-Stat3^-/-^* chimera). Circles in color represent significant differentially expressed (DE) genes. Highlighted are selected genes of the pathway “Fisher G2/M cell cycle”. **(B**) Pathway analysis of all significant DE genes from A). **(C - D)** Cell Trace Violet (CTV) dilution assay of WT and *NK-Stat3^-/-^* NK cells stimulated with a titration of IL-15 in the presence of absence of IL-10, with representative CTV histograms shown. **(E)** Schematic for RNA-seq experiment of *in vitro* stimulated NK cells. **(F)** Volcano plot of NK cells stimulated with IL-15 versus IL-10 + IL-15. Significant DE genes are shown in blue, with outlines highlighting transcription factors, and gene names highlighted from a cell proliferation pathway (see methods). **(G)** Quantification of up- or downregulated gene number in conditions from (F) and Fig. S3A. Values indicate (# upregulated DE genes) / (# downregulated DE genes). **(H)** Scatterplot of expected versus observed log2 fold change in RNA-seq experiment. Circles in color are genes which the absolute difference between the observed and expected log2 fold change values exceed 0.8 and which is significantly DE in any of the two individual contrasts. Genes of interest are labelled. Data in 2C and 2D are representative of two independent experiments.

To mechanistically investigate the impact of STAT3 following IL-15 signaling, we performed RNA-seq of NK cells stimulated with IL-10, IL-15, or a combination of both (**Fig. 2E** and **Supp. Fig. 3A**). Individually, each cytokine induced a distinct transcriptional pattern: IL-15 stimulation upregulated as many genes as were downregulated, whereas IL-10 regulated far fewer genes overall but globally induced rather than suppressed gene transcription (**Supp. Fig. 3A**). Addition of IL-10 modulated IL-15 signaling and resulted in an overall increase in induced over repressed genes (**Fig. 2F**-**G**, **Supp. Fig. 3B**) We hypothesized that IL-10 and IL-15 could synergize to promote NK cell proliferation in one of two ways: 1) IL-10 and IL-15 could separately induce transcriptional changes via STAT3 and STAT5, respectively, which together optimally promote NK cell proliferation. 2) STAT3 and STAT5 activation via IL-10 and IL-15, respectively, could indirectly affect each other, possibly via an epigenetic mechanism. By plotting expected (log2 fold change [IL-10 versus Unstim] + log2 fold change [IL-15 versus Unstim]) against observed log2 fold change (log2 fold change [IL-10 + IL-15 versus Unstim]), we found evidence for both mechanisms. Although many genes were regulated in a manner predicted by the two isolated stimuli, other genes did not show such a pattern (**Fig. 2H**), genes outside of the dotted line borders). Together, we observe that STAT3 activation via IL-10 will alter IL-15 signaling through direct and indirect mechanisms to promote proliferation of NK cells under homeostatic conditions.

### STAT3 competes with STAT5 to regulate *Myb*, a negative regulator of NK cell differentiation

To further dissect how STAT3 modulates IL-15 signaling in NK cells, we performed STAT3 and STAT5 CUT&RUN on NK cells stimulated with IL-10 (canonical STAT3 activator), IL-15 (canonical STAT5 activator) or IL-10 + IL-15 (**Fig. 3A-B**). Interestingly, addition of IL-10 to IL-15 overall reduced STAT5 genomic binding (**Fig. 3A**), while more STAT3 binding was observed in IL-10 + 15 than IL-10 alone (**Fig. 3B**). We hypothesized that STAT3 competes with STAT5 for genomic binding. Indeed, many regions showing IL-15 dependent STAT3 binding simultaneously showed less STAT5 binding (**Fig. 3C-D, Supp Fig. 3C**).

**Figure 3:**
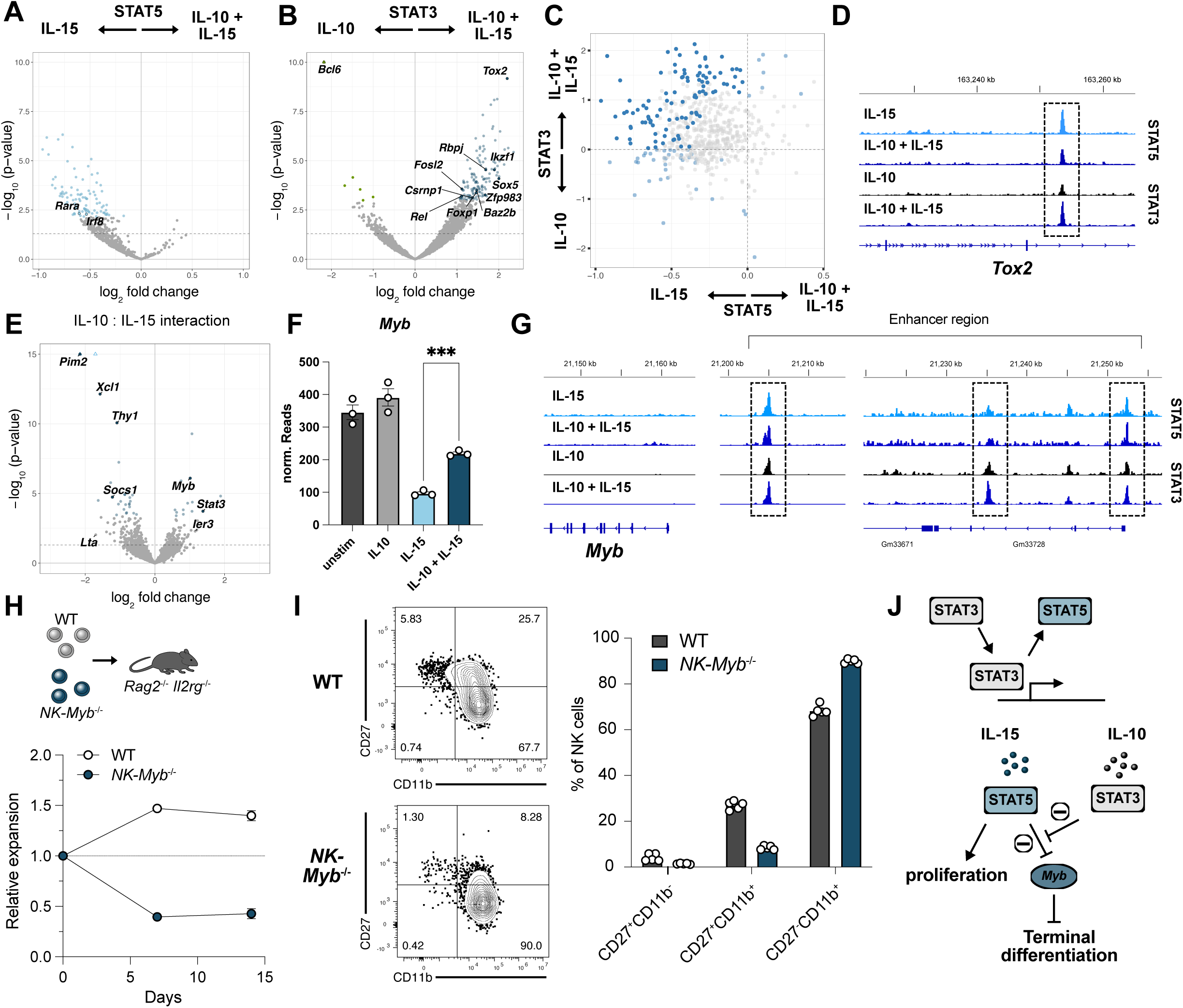
Competition of STAT3 and STAT5 for genomic binding regulates *Myb*, a negative regulator of NK cell differentiation. **(A)** Volcano plot of differential binding (DB) analysis from STAT5 CUT&RUN in NK cells stimulated with IL-15 versus IL-10 + IL-15. Blue circles highlight significantly DB regions in transcription factor loci. **(B)** Volcano plot of STAT3 CUT&RUN shows DB analysis in NK cells stimulated with IL-10 versus IL-10 + IL-15. Blue circles highlight significantly DB regions in transcription factor loci. **(C)** Scatterplot of data from (A) and (B) for regions commonly bound by STAT3 and STAT5, with significant DB peaks shown in blue. **(D)** Representative tracks are shown for the *Tox2* gene. **(E)** Interaction analysis of RNA-seq experiment in Fig. 2F. Genes showing significant interaction reveal an expression pattern in IL-10 + IL-15 stimulated NK cells not anticipated by IL-10 stimulation alone. Blue circles are genes with significant interactions, and highlighted circles are select genes of interest. **(F)** Normalized Reads of *Myb* gene expression showing IL-10 only increases *Myb* significantly in the presence of IL-15. **(G)** Representative tracks in *Myb* enhancer peaks for STAT3 and STAT5 CUT&RUN as shown in (A) and (B). **(H)** Transfer of WT and *NK-Myb^-/-^* NK cells into *Rag2^-/-^ x Il2rg2^-/-^* mice and analysis of blood at various time points shown. **(I)** Phenotype (CD27 and CD11b subsets^56^) of WT and *NK-Myb^-/-^*NK cells at steady state. **(J)** Model shows IL-15 suppression of *Myb* expression via STAT5 leading to terminal differentiation, with STAT3 preventing this STAT5-induced terminal differentiation. Data in 3H is representative of 2 independent experiments. Data in 3I is representative of 3 individual experiments. Significance in 3F was calculated using paired t-test (same biological replicates): *** p < 0.001, ** p < 0.01, * p < 0.05.

A previous report suggested that STAT3 and STAT5 can have opposite effects on gene expression^29^. We proposed that replacement of STAT5 with STAT3 could modulate expression of downstream transcription factors crucial for NK cell proliferation. To identify candidates that show evidence of such antagonistic transcriptional regulation, we performed an interaction analysis of our RNA-seq dataset (**Fig. 3E**). Interaction analysis analyzes how the degree of differences in expression observed after IL-10 stimulation are modulated by IL-15. Among the few TFs that displayed an antagonistic regulation we found *Myb*, a transcription factor that can prevent terminal differentiation in CD8^+^ T cells^30^. Addition of IL-10 antagonized IL-15 dependent suppression of *Myb* expression, even though IL-10 stimulation alone did not substantially induce *Myb* transcription (**Fig. 3F**). In CUT&RUN analysis, a known enhancer region of *Myb*^31^ showed increased STAT3 genomic binding after stimulation with IL-10 + IL-15 compared to IL-10 alone and a concomitant reduction in STAT5 binding at certain sites following STAT3 activation (**Fig. 3G**).

Because MYB may act as a key downstream transcription factor regulated by STAT3 under homeostatic conditions, we tested if MYB affects homeostatic NK cell expansion similar to STAT3 (**Fig. 1H**) using a model of conditional MYB deficiency (*Nkp46^Cre^ x Myb^flox/flox^*). Indeed, *NK-Myb^-/-^*NK cells showed an expansion defect similar to *NK-Stat3^-/-^* NK cells in *Rag2^-/-^ x Il2rg^-/-^* mice (**Fig. 3H and 1H**). Furthermore, *NK-Myb^-/-^* NK cells showed a more mature phenotype under steady state (**Fig. 3I**), suggesting that like in CD8^+^ T cells^30^, *Myb* may block terminal differentiation in NK cells. In summary, we propose that STAT3 antagonizes STAT5 dependent suppression of genes by directly competing with STAT5 for genomic binding. This antagonism includes *Myb*, which is downregulated by IL-15 without simultaneous activation of STAT3. In the absence of STAT3 activation, our findings suggest IL-15 drives terminal differentiation via suppression of *Myb*, leading to suboptimal expansion potential (**Fig. 3J**).

### STAT3 modulates type-I IFN signaling in antiviral NK cells during high-dose infection

Under homeostatic conditions, we observed that STAT3 interacts with IL-15 signaling to promote proliferation by modulating downstream transcription factor networks including MYB. Because of the similarity between STAT3-dependent TF expression under homeostasis and low-dose infection (**Fig. 1G**), we hypothesized that such a mechanism in part explains the positive impact of STAT3 on NK cell proliferation in low-dose infection. However, why does STAT3 in high-dose infection show an opposite and detrimental effect on adaptive NK cell responses?

Comparing transcriptomes of WT NK cells in low- versus high-dose infection (**Fig. 4A**), we found an enrichment in pathways related to type-I IFN signaling (**Fig. 4B** and **1B**). This enhanced interferon signaling depends in part on STAT3, as these pathways were only significantly enriched in WT but not *NK-Stat3^-/-^* NK cells during high-dose infection (**Fig. 4B-C**). To substantiate the hypothesis that high-dose infection is characterized by enhanced interferon signaling in NK cells, we compared high-dose dependent genes to datasets previously generated using STAT- and cytokine receptor-deficiency models^10,32–35^ (**Fig. 4D** and **Supp. Fig. 4A-F**). Only STAT1- and IFNAR-dependent genes showed a positive correlation with high-dose dependent genes. At the protein level, type-I IFN and IFN-γ, the latter produced by NK cells, were robustly elevated in high-dose infection (**Fig. 4E**, top). At the same time, other cytokines such as IL-10 and IL-12 were only slightly elevated, and IL-15 levels remained constant between low- and high-dose infection (**Fig. 4E**, bottom). These data together suggest that high-dose infection specifically leads to elevated interferon levels, and concomitant interferon signaling in NK cells modulated by STAT3.

**Figure 4:**
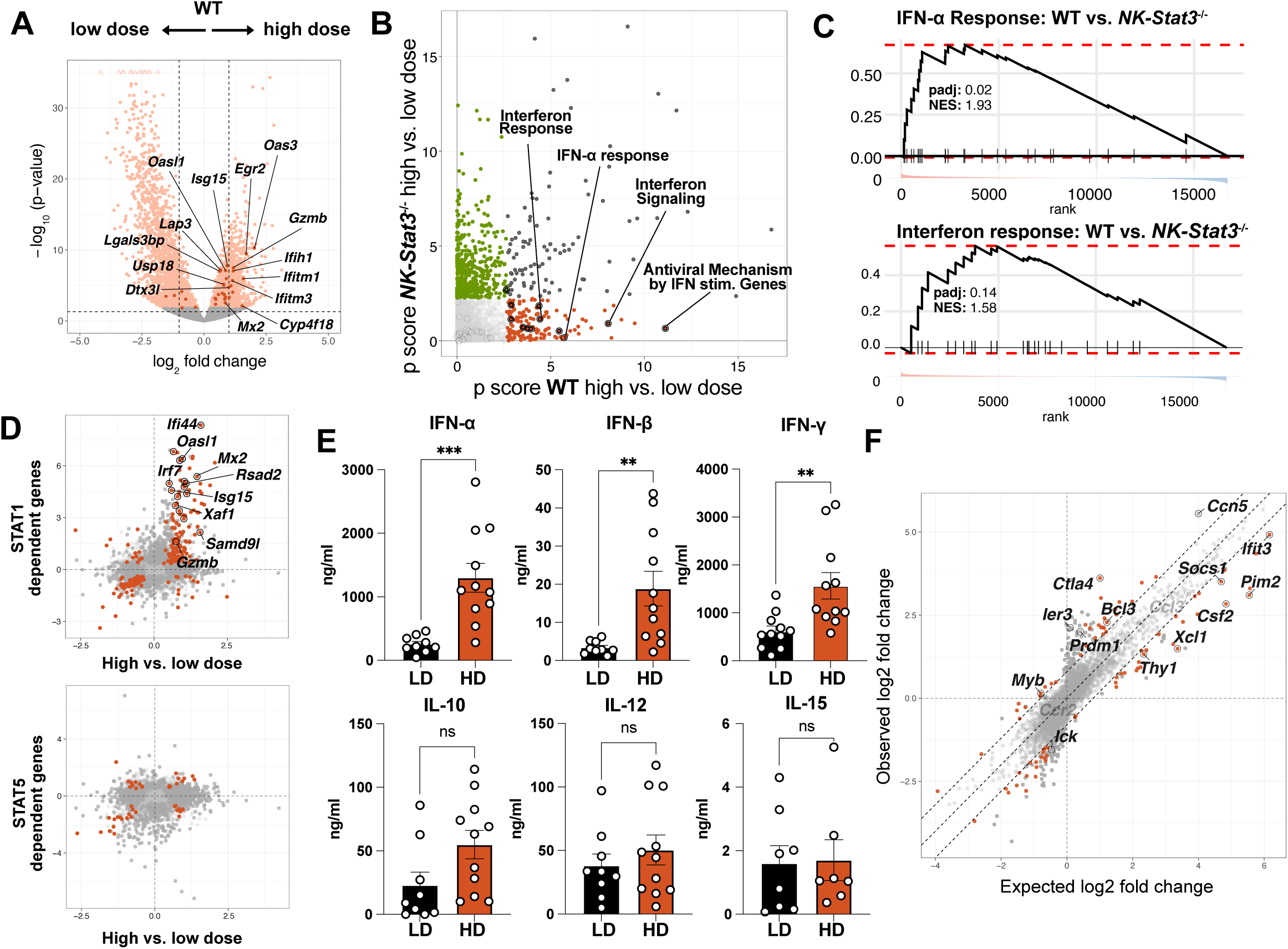
High-dose MCMV infection is characterized by STAT3-dependent type-I interferon signaling in NK cells. **(A)** Volcano plot of RNA-seq analysis in WT NK cells during HD and LD infection at day 4 PI. Circles in color are significantly differentially expressed genes and highlighted genes are from the enriched IFN-α Response pathway in (B). **(B)** GO Pathway analysis for WT and *NK-Stat3^-/-^* NK cells in HD versus LD infection. ‘p score’ indicates -log10(p-value). Circles in color indicate significant pathway (padj ≤ 0.05). Black circles indicate interferon associated pathways. **(C)** GSEA plot of two of the interferon pathways shown in (B). **(D)** Scatterplot of differentially expressed genes in HD versus LD infection (x-axis) compared with genes differentially expressed in WT versus *Stat1^-/-^* or *NK^Cre^ ^x^ Stat5^flox/wt^* mice (y-axis) during MCMV infection. Highlighted are genes from the “Interferon response” pathway. **(E)** Serum levels of cytokines on day 2 of LD versus HD infected mice. **(F)** Scatterplot of expected versus observed log2 fold change of IL-10 and IFN-α as described in Fig. 2H. Data in 4E is pooled from 2 individual experiments. Significance in 4E was calculated using unpaired t-test: *** p < 0.001, ** p < 0.01, * p < 0.05.

To test whether activation of STAT3 via IL-10 can modulate interferon signaling in NK cells as observed for IL-15, we performed RNA-seq of NK cells stimulated with IL-10, IFN-α or a combination of both (**Supp. Fig. 5A**). As we observed for IL-15, IFN-α stimulation altered expression of far more genes and was less biased towards gene induction than IL-10 stimulation (**Supp. Fig. 5B**). To discern direct from indirect interactions on transcription we again plotted expected versus observed changes for co-stimulation with IL-10 and IFN-α (**Fig. 4F**). We observed an even higher number of genes that showed evidence of an indirect regulation, meaning that the observed changes did not fit the prediction from adding the log2 fold changes of either stimulus alone. Thus, these findings suggest STAT3 modulates type-I-IFN signaling in NK cells during high-dose infection, in part via indirect mechanisms of action.

### Inflammation-dependent STAT3 redirection instructs a TF network including *Prdm1*/BLIMP-1 that promotes terminal differentiation

To better understand the mechanism of STAT3-dependent modulation during inflammatory cytokine signaling, we performed CUT&RUN for STAT1 and STAT3 following stimulation of primary NK cells with different cytokines. Because MCMV infection is characterized by high levels of both interferon and IL-12, we included this proinflammatory cytokine to more optimally mimic inflammation during infection. In contrast to our observations for STAT5, activation of STAT3 via IL-10 did not significantly alter STAT1 binding following proinflammatory IFN-α + IL-12 treatment (**Fig. 5A**). In contrast, IFN-α + IL-12 dramatically modulated STAT3 genomic binding to key TFs (**Fig. 5B**). Although no significant reduction in STAT1 binding following STAT3 activation was observed, STAT3 binding sites showed a bias towards lower STAT1 binding (**Fig 5C**), indicating that competition may represent a mechanism contributing to STAT relocation. To identify the most crucial TFs regulated by STAT3 in the context of interferon signaling, we performed an interaction analysis from RNA-seq data to assess how the degree of differences driven by IL-10 are modified in presence of IFN-α (**Fig. 5D**). One of the most altered TFs in this analysis was *Prdm1*, encoding for the protein BLIMP-1. Only a combination of IL-10 + IFN-α, but not each cytokine alone, induced *Prdm1* expression (**Fig. 5E**). Several regions within and flanking the *Prdm1* gene showed a robust increase in STAT3 binding under inflammatory conditions (**Fig. 5F-G**). Thus, STAT3 regulates *Prdm1* expression in the presence of inflammatory cytokines through preferential and direct binding to the *Prdm1* locus.

**Figure 5:**
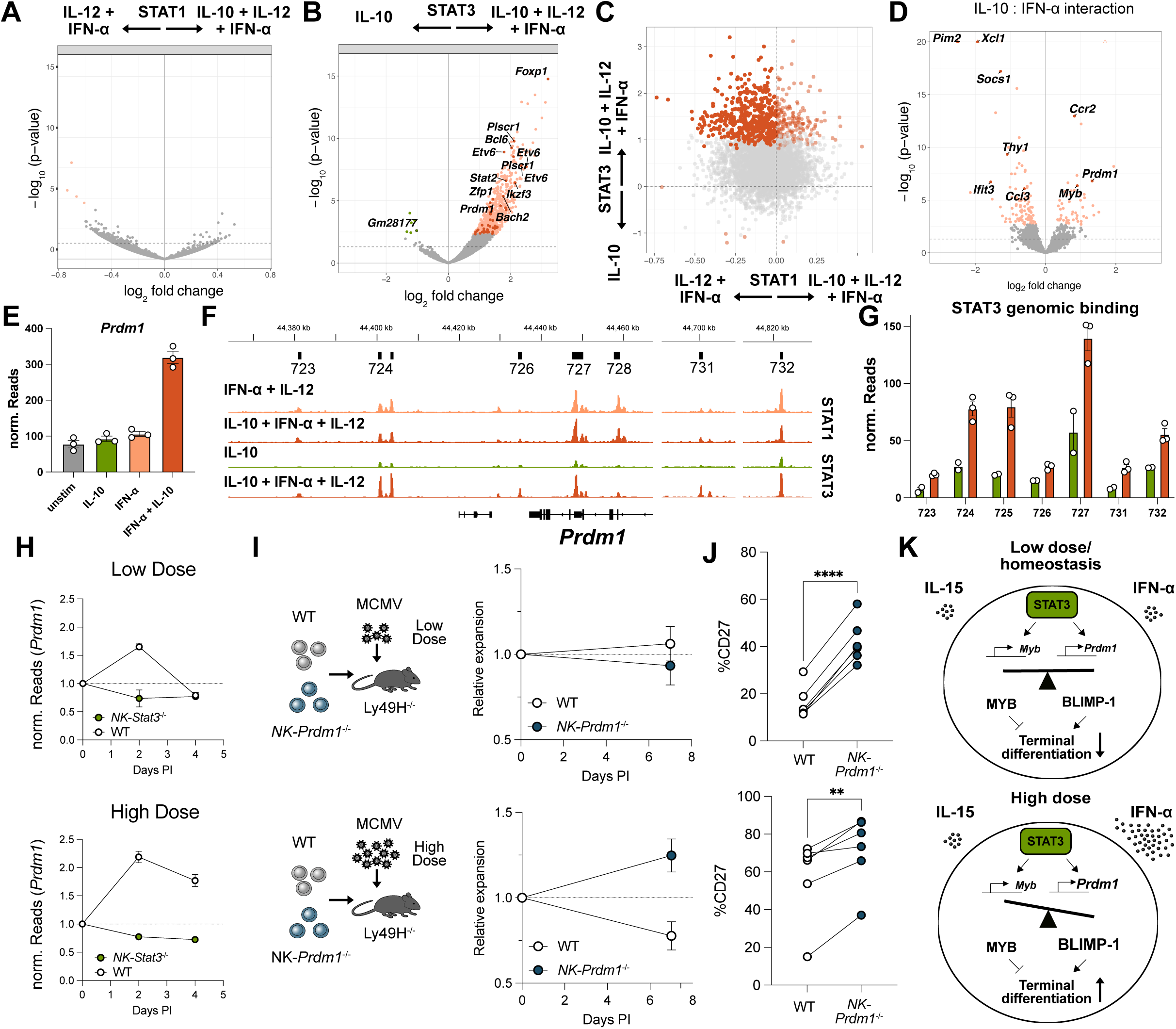
Interferon-dependent relocation of STAT3 regulates *Prdm1*/BLIMP-1 to promote NK terminal differentiation during high-dose infection. **(A)** Volcano plot shows DB analysis of STAT1 CUT&RUN in NK cells stimulated with IL-10 + IL-12 + IFN-α versus IL-12 + IFN-α. **(B)** Volcano plot shows DB analysis of STAT3 CUT&RUN in NK cells stimulated with IL-10 + IL-12 + IFN-α versus IL-10. Red circles are significantly DB regions with specific TFs highlighted. **(C)** Scatterplot of data from (A) and (B) for regions commonly bound by STAT3 and STAT1. Red circles are significant DB peaks. **(D)** Interaction analysis of RNA-seq experiment in Fig. 4F shown. Genes showing significant interaction reveal an expression pattern in IL-10 + IFN-α stimulated NK cells not anticipated by IL-10 stimulation alone. Red circles are genes with significant interactions, and highlighted circles are select genes of interest. **(E)** Normalized Reads of *Prdm1* gene expression shown for NK cells stimulated with IL-10, IFN-α, or a combination of IL-10 + IFN-α. **(F)** Representative tracks show STAT1 and STAT3 CUT&RUN within and flanking the *Prdm1* gene. **(G)** Quantification of STAT3 peaks shown in (F). **(H)** Normalized *Prdm1* reads (relative to day 0) in WT and *NK-Stat3^-/-^* NK cells during LD or HD infection with MCMV at various time points. **(I)** Graphs show WT and *NK-Prdm1^-/-^* NK cells co-transferred into *Ly49h^-/-^* mice infected with LD (top) or HD (bottom) of MCMV. **(J)** CD27 expression in NK cell populations shown in (I). **(K)** Model highlights STAT3 operating as a transcriptional switch, regulating downstream transcription factors *Myb* and *Prdm1*/BLIMP-1 in homeostatic versus inflammatory environments. Data in 5I - 5J is pooled from 2 individual experiments. Significance in 5J was calculated using paired t-test (same biological replicates due to co-transfer): *** p < 0.001, ** p < 0.01, * p < 0.05.

We hypothesized that this inflammation-dependent regulation of *Prdm1* may contribute to the context-dependent role of STAT3 observed during MCMV infection (**Fig. 1C-E**). Indeed, WT NK cells upregulated *Prdm1* expression as early as day 2 PI and showed elevated and prolonged expression during high-dose infection (**Fig. 5H**). In contrast, STAT3-deficient NK cells showed no evidence of such regulation (**Fig. 5H**). To explain the context-dependent phenotype of STAT3-deficient NK cells, we hypothesized that BLIMP-1 inhibits NK cell expansion specifically in high-dose infection. As we have observed previously^36^, deletion of *Prdm1* in NK cells did not impact their expansion following low-dose infection; however, in high-dose infection, Prdm1-deficient NK cells (*Nkp46^Cre^ x Prdm1^flox/flox^*) outcompeted WT counterparts (**Fig. 5I**). Furthermore, *NK-Prdm1^-/-^* NK cells exhibited a less differentiated CD27^+^ phenotype following both low- and high dose infection (**Fig. 5J**). These findings suggest that, in high-dose infection, STAT3 induces *Prdm1*/BLIMP-1 drive terminal maturation that inhibits NK cell expansion. Altogether, we have uncovered a novel inflammation sensory mechanism by which context-dependent relocation of STAT3 establishes STAT3 as a transcriptional switch, instructing different downstream transcription factors including MYB and BLIMP-1, with opposing roles in adaptive potential versus terminal differentiation dependent on the presence of homeostatic versus inflammatory cytokines (**Fig. 5K**).

## DISCUSSION

Cytokine signaling via STAT transcription factors is one of the key pathways instructing immune cell fate. However, despite many decades of research the precise mechanisms underlying how STATs shape cell differentiation and function remain incompletely understood. Complexity on different levels of signal integration contributes to a context-dependent role for both cytokines and STAT transcription factors, including the redundant activation of JAK adapters^23^, the formation of STAT homo- and heterodimers^37^, and the epigenetic interactions of STATs^38,39^. In this study, we find that STAT3 is relocated to distinct genomic sites under homeostatic or inflammatory conditions, differentially mediated by IL-15 or type-I IFN signaling, respectively. The development of our low-input CUT&RUN enables such an analysis, which bypasses the complex upstream mechanisms of cytokine signal integration to allow direct visualization of differential STAT genomic binding. However, dissecting the precise upstream mechanisms contributing to STAT genomic relocation will be investigated in the future.

Our observation poses a crucial question for the STAT3 inhibitors currently being tested in immunotherapy against cancer: does the impact of STAT3 inhibitors depend on the level of inflammation encountered in tumors? Solid cancer can be grouped into “hot” versus “cold” tumors based on the status of immune activation^40^. Our data predicts that in cold tumors STAT3 inhibitors may repress while in hot tumors they would enhance NK cell proliferation. Because clinical responses to STAT3 inhibitors are highly variable^41^, adapting the use of these inhibitors to the level of tumor immune activation, or combination of these inhibitors with treatments that increase tumor inflammation, may be promising strategies to improve tumor therapy.

Understanding the dual and often opposite role played by STAT3 we observe in this study will be crucial beyond NK cell differentiation and function. For instance, STAT3 gain-of-function mutations (STAT3^GOF^) are common in NK cell lymphoma^42^, where malignant cells will acquire mutations that promote stemness and antagonize terminal differentiation. Our study predicts that STAT3 activation could therefore have both positive and negative roles for lymphoma progression. Under homeostatic conditions, STAT3 would upregulate *Myb*, a known driver of lymphoma stemness and proliferation^43^, whereas during inflammation *Prdm-1* induction may lead to terminal differentiation of STAT3^GOF^ lymphomas. Interestingly, a recent study found that STAT3^GOF^ mutations are often accompanied by Prdm-1 loss-of-function mutations in NK cell lymphoma, fitting predictions from our study^44^. Because many hematologic and solid cancers show increased STAT3 activity and/or STAT3^GOF^ mutations^45,46^, this observation suggests an intriguing therapeutic concept: is it possible to deviate STAT3 genomic binding from “adaptive or stemness” programs towards promoting terminal differentiation in cancer cells? A similar strategy targeting a pathway promoting differentiation of myeloid cells is currently being employed in treatment of acute promyelocytic leukemia using all-trans retinoic acid (ATRA)^47^.

Beyond STAT3, context-dependent roles have been described for additional cytokines. For example, in T cells, STAT5 can exert different functions downstream of IL-2, IL-7 and IL-15 receptors^48–51^. Futhermore, STAT1 downstream of interferons can support memory and effector^5,52,53^ differentiation, but also drive exhaustion^54,55^. The approaches in our study offer a practical means to dissect the underlying mechanisms of such context-dependent cytokine effects in primary cells. These advances not only deepen our mechanistic understanding of cytokine responses but may also pave the way for new therapeutic strategies that target context-specific signaling in human disease.

## Acknowledgments

We thank past and present members of the Sun lab for helpful discussions and support. Sequencing was performed by the Integrated Genomics Operation Core which is funded by the NCI Cancer Center Support Grant (CCSG, P30 CA08748), Cycle for Survival, and the Marie-Josée and Henry R. Kravis Center for Molecular Oncology. S.G. is the recipient of a CRI / Donald J. Gogel Postdoctoral Fellowship (CRI Award CR13934), was supported by the Deutsche Forschungsgemeinschaft (DFG, Award number GR5503/1-1) and NIAID (NIH) under award number K99AI180360. S.X.F. was supported by a Medical Scientist Training Program grant from the National Institute of General Medical Sciences of the National Institutes of Health under award number T32GM152349 to the Weill Cornell/Rockefeller/Sloan Kettering Tri-Institutional MD-PhD Program. J.C.S. was supported by the Ludwig Center for Cancer Immunotherapy, the American Cancer Society, the Burroughs Wellcome Fund, and the NIH (AI100874, AI130043, AI155558, and P30CA008748).

## MATERIALS AND METHODS

### Mice

All mice used in this study were housed and bred under specific-pathogen-free conditions with food and water in 12-h light–dark cycles at 72 °F with 30–70% humidity at Memorial Sloan Kettering Cancer Center and handled in accordance with the guidelines of the Institutional Animal Care and Use Committee. The following mouse strains were used in this study: C57BL/6 (CD45.2), C57BL/6 CD45.1 (CD45.1), C57BL/6 CD45.1×CD45.2, Ncr1-Cre x STAT3^flox/flox^ (*NK-Stat3^−/−^*), Ncr1-Cre x *Myb*^flox/flox^ (*NK-Myb^−/−^*), Ncr1-Cre x Prdm1^flox/flox^ (*NK-Prdm1^−/−^*), Klra8^−/−^ CD45.1xCD45.2 (Ly49H-deficient). Experiments were conducted using 8–10-week-old mice or 8–16 weeks post-transplant mixed bone-marrow chimeric mice and all experiments were conducted using age- and sex-matched mice in accordance with approved institutional protocols.

### MCMV virus preparation

MCMV (Smith strain) was serially passaged through BALB/c hosts three times and then salivary gland viral stocks were prepared with a homogenizer for dissociating the salivary glands of infected mice 3 weeks after infection.

### Mixed bone-marrow chimeras

Mixed bone-marrow chimeric (mBMC) mice were generated by lethally irradiating (950LcGy) C57BL/6 CD45.1×CD45.2 animals and reconstituting with a 1:1 mixture of bone-marrow cells from WT (CD45.1) and *Il12rb2^−/−^* (CD45.2) mice. Hosts were co-injected with anti-NK1.1 (PK136) to deplete any residual donor and host NK cells. Residual CD45.1^+^CD45.2^+^ host NK cells were excluded from all analyses. Chimerism in WT:*NK-Stat3^-/-^* chimera was assessed by blood staining 1-2 weeks prior to the experiment.

### Infections

Mice were infected i.p. with 1 x 10^3^ PFU (low-dose) or 5 x 10^3^ PFU (high-dose) of MCMV and analyzed at the respective time points using flow cytometry.

### Isolation of mouse NK cells and flow cytometry

Spleens were dissociated with glass slides and filtered through a 100-μm cell strainer. Flow cytometry experiments were analyzed using a Cytek Aurora (Cytek Biosciences). Cell sorting was performed using a BD Aria II cytometers (BD Biosciences). Before cell sorting, NK cells were enriched by incubating whole splenocytes with the following antibodies at 10 μg/ml: CD3ε (Clone 17A2), CD4 (Clone GK1.5), CD8 (Clone 2.43), Ter119 (Clone TER-119), CD19 (Clone 1D3), Ly6G (Clone 1A8) (BioXCell). After washing, cells were incubated with goat anti-rat beads (QIAGEN, cat. no. 310107). For intracellular staining, cells were fixed and permeabilized using eBioscience Intracellular Fix & Perm Buffer Set (Thermo Fisher, cat. no. 88-8824-00).

### Cell culture for *in vitro* experiments

NK cells were cultured in complete IMDM (10% FBS, 1× L-glutamine, 1× sodium pyruvate, 1× BME, 1× MEM-NAA and 25 mM HEPES). For cytokine stimulations, we used 50 ng/ml mouse IL-15 (Peprotech, cat. no. 210-15), 1 x 10^2^ U/ml mouse IFN-α (R&D Cat. 12105-1), 20 ng/ml mouse IL-12 (R&D Systems, cat. no. 419-ML-050), and 10 ng/ml mouse IL-10 (Peptrotech), unless otherwise indicated (e.g. Fig. 2C).

### Cytokine stimulation for sequencing

Cytokine stimulations were performed for 1h (CUT&RUN experiments) or 3h (RNA-seq experiments) in complete IMDM (10% FBS, 1× L-glutamine, 1× sodium pyruvate, 1× BME, 1× MEM-NAA and 25 mM HEPES).

### CTV dilution and proliferation assay

Splenocytes from CD45.1/.1 (WT1) and *NK-Stat3*^-/-^ mice were pooled and enriched using the NK Cell Isolation Kit (Miltenyi, Cat. 130-115-818). For CTV assays, cells were stained with CellTrace Violet (CTV) according to the manufacturer’s instructions (Thermo Scientific, cat. C34557).

### Serum protein quantification

Serum was collected by coagulating blood for 30 minutes at room temperature and centrifugation at 2000g for 15min. IL-10, IL-12, IFN-α, IFN-β, and IFN-γ were assessed using a Legendplex Custom panel (Biolegend). IL-15 was quantified using Mouse IL-15 ELISA Development Kit (ABTS) (Thermo).

### Illumina Sequencing of RNA-seq from high-dose and low-dose infection

RNA was extracted using Arcturus PicoPure RNA Isolation Kit (Cat. 12204-01) After RiboGreen quantification and quality control by Agilent BioAnalyzer, 0.45-1.5 ng total RNA with RNA integrity numbers ranging from 8.5 to 10 underwent amplification using the SMART-Seq v4 Ultra Low Input RNA Kit (Clonetech catalog # 63488), with 12 cycles of amplification. Subsequently, 8.5 ng of amplified cDNA was used to prepare libraries with the KAPA Hyper Prep Kit (Roche 07962363001) using 8 cycles of PCR. Samples were barcoded and run on a NovaSeq 6000 in a PE100 run, using the NovaSeq 6000 S4 Reagent Kit (200 cycles) (Illumina). An average of 40 million paired reads were generated per sample, the percent of mRNA bases per sample ranged from 75% to 87%, and ribosomal reads averaged 0.22%.

### Mercurius BRB-seq

RNA was extracted using ZymoResearch Quick RNA MicroPrep Kit. RNA samples were multiplexed in 96 well plates containing oligos and processed according to the manufacturer’s instructions (MERCURIUS™ Standard BRB-seq library preparation kit, cat 11013).

### Transcription factor CUT&RUN

For STAT1, STAT3, and STAT5 CUT&RUN, 300,000-500,000 sorted total NK cells were used. Cells were light fixated with 0.1% PFA and fixated in antibody buffer (1X eBioscience Perm/Wash Buffer, 1X Roche cOmplete EDTA-free Protease Inhibitor, 0.5 uM Spermidine, + 2uM EDTA in H_2_O) prior to “staining” with antibodies at a 1:100 dilution: STAT1 (Proteintech Cat. 10144-2-AP), STAT3 (CellSignaling Clone 124H6, Cat. 9139) and STAT5 (Thermo, Cat. 71-2500). Upon antibody incubation, cells were washed twice with Buffer 1 (1X eBioscience Perm/Wash Buffer, 1X Roche cOmplete EDTA-free Protease Inhibitor, 0.5 uM Spermidine in H_2_O) and resuspended in 50ul of Buffer 1 + 1X pA/G-MNase (Cell Signaling, cat. 57813) and incubated for 1 hour at 4°C. Cells were washed with Buffer 2 (0.05% w/v Saponin, 1X Roche cOmplete EDTA-free Protease Inhibitor, 0.5 uM Spermidine in 1X PBS) three times. After washing, Calcium Buffer (Buffer 2 + 2uM of CaCl_2_) was used to resuspend the cells for 30 mins at 4°C to activate the pA/G-MNase reaction, and equal volume of 2X STOP Buffer (Buffer 2 + 20uM EDTA + 4uM EGTA) was added along with 1 pg of *Saccharomyces cerevisiae* spike-in DNA (Cell Signaling, cat. 29987). Samples were incubated for 15 mins at 37°C and DNA was isolated and purified using Qiagen MinElute Kit according to manufacturer’s protocol and subjected to library amplification^57,58^.

### Illumina Sequencing of CUT&RUN and Mercurius libraries

After PicoGreen (CUT&RUN) quantification and quality control by Agilent TapeStation, the libraries were pooled equimolar and sequenced on a NovaSeq 6000 in a PE100 run, using the NovaSeq 6000 S4 Reagent Kit (200-300 Cycles) (Illumina).

### RNA-seq and CUT&RUN processing and analysis

Data processing methods for published RNA-seq datasets have been previously described^33,35^. For both RNA-seq datasets, transcript annotation was based on the mm10 University of California, Santa Cruz (UCSC) Known Gene model. For dosage RNA-seq dataset generated in this study, transcript quantification was performed using the quasi-mapping-based mode of Salmon (v0.13.1) correcting for potential GC bias. Transcript was summarized to gene level using tximport (v1.10.1). For cytokine RNA-seq dataset generated in this study, transcript alignment was performed using STAR (v2.7.11b_alpha_2024-02-09) as described in manufacturer’s BRB-seq protocol (Alithea MERCURIUS kit, rev Jan 2025). Reads assigned to transcripts were summed to gene level using transcript to gene information in TxDb.Mmusculus.UCSC.mm10.knownGene (v3.10). For CUT&RUN datasets found in this study, paired reads were trimmed for adaptors and removal of low-quality reads by using Trimmomatic (v0.39) and aligned to the mm10 reference genome using Bowtie 2 (v2.4.1). Upon alignment, peaks were called using MACS2 (v2.2.7.1) with input samples as a control using narrow peak parameters with cutoff-analysis -p 1e-5 --keep-dup all -B -- SPMR. For each CUT&RUN targets, a peak atlas was generated by merging via union of peaks called by each sample after filtering out bottom 25% of peaks by MACS2 calculated qValue. Resulting peak atlas was further filtered to only retain peaks that were called by two or more replicates within each sample group, as well as peaks that do not overlap the Encode’s DAC Exclusion List Regions (ENCFF547MET). Reads were mapped to the respective final atlas and counted with the summarizeOverlaps function from the GenomicAlignment package (v1.34.1). CUT&RUN data was normalized using size factor values calculated from aligned *Saccharomyces cerevisiae* spike-in DNA counts through DESeq2 (v1.46.0) ‘estimateSizeFactors’ function. For data generated in this study, differential analyses were executed with DESeq2 using UCSC Known Gene models as reference annotations. For RNA-seq and CUT&RUN, genes/peaks were considered differential if they showed adjusted p-value (padj) ≤ 0.05. For tracks, BAM files were converted to bigwig files using bamCoverage function and scaled based on size factor as described above. Scaled bigwig files from each replicate per condition were then averaged using bigwigAverage function from deepTools (v3.5.4).

### Pathway analysis

Gene ontology analysis used goseq R package (v1.58.0) and GSEA analysis used fgsea R package (v1.32.0) with MSigDB gene sets C2 and C5. The names of MSigDB pathways are abbreviated as follows: Cell Cycle = Fischer G2 M cell cycle, IFN-α Response = Moserle IFNa Response, Mitotic Spindle Checkpoint = Reactome Mitotic Spindle Checkpoint, STAT3 Targets = Dauer STAT3 Targets DN, IFNa/b signaling = Reactome Interferon Alpha Beta Signaling, Myc targets = Yu Myc Targets Up, Cycling Genes = Benporath Cycling Genes, APC/MYC targets = Sansom APC MYC targets, MYB Family targets = Lang MYB Family Targets, Cell Cycle = Fisher G2/M Cell Cycle, IL2 pathway = PID IL2 PATHWAY, Interferon response = Zhang Interferon response, IFN-α response = Moserle IFNa response, Interferon signaling = Reactome interferon signaling, Antiviral mechanism by IFN stim. genes = Reactome Antiviral mechanism by IFN stimulated genes. The cell proliferation pathway highlighted in Fig. 2F represents the C5 GO BP term “REGULATION OF CELL POPULATION PROLIFERATION”.

### Expected versus observed and interaction analysis

For expected versus observed analysis, expected value was calculated based on the addition of log2 fold change values of individual stimuli versus control: log2 fold change [stim A versus unstimulated] + log2 fold change [stim B versus unstimulated], where A and B are either IL-10 and IL-15, or IL-10 and IFN-α. Observed value was calculated based on the direct contrast of log2 fold change [stim A + stim B versus unstimulated]. A gene was deemed significant based on statistics (padj ≤ 0.05) in either of the two individual contrasts used for expected value. Interaction analysis used “∼ stim A + stim B + stim A:stim B” (IL-10 and IL-15, or IL-10 and IFN-α) as GLM design in DESeq2. The results from the interaction term contrast (stim A:stim B) was used and a gene was deemed to have significant interaction based on statistics (padj ≤ 0.05).

## SUPPLEMENTARY FIGURE LEGENDS

**Supp. Fig. 1:**
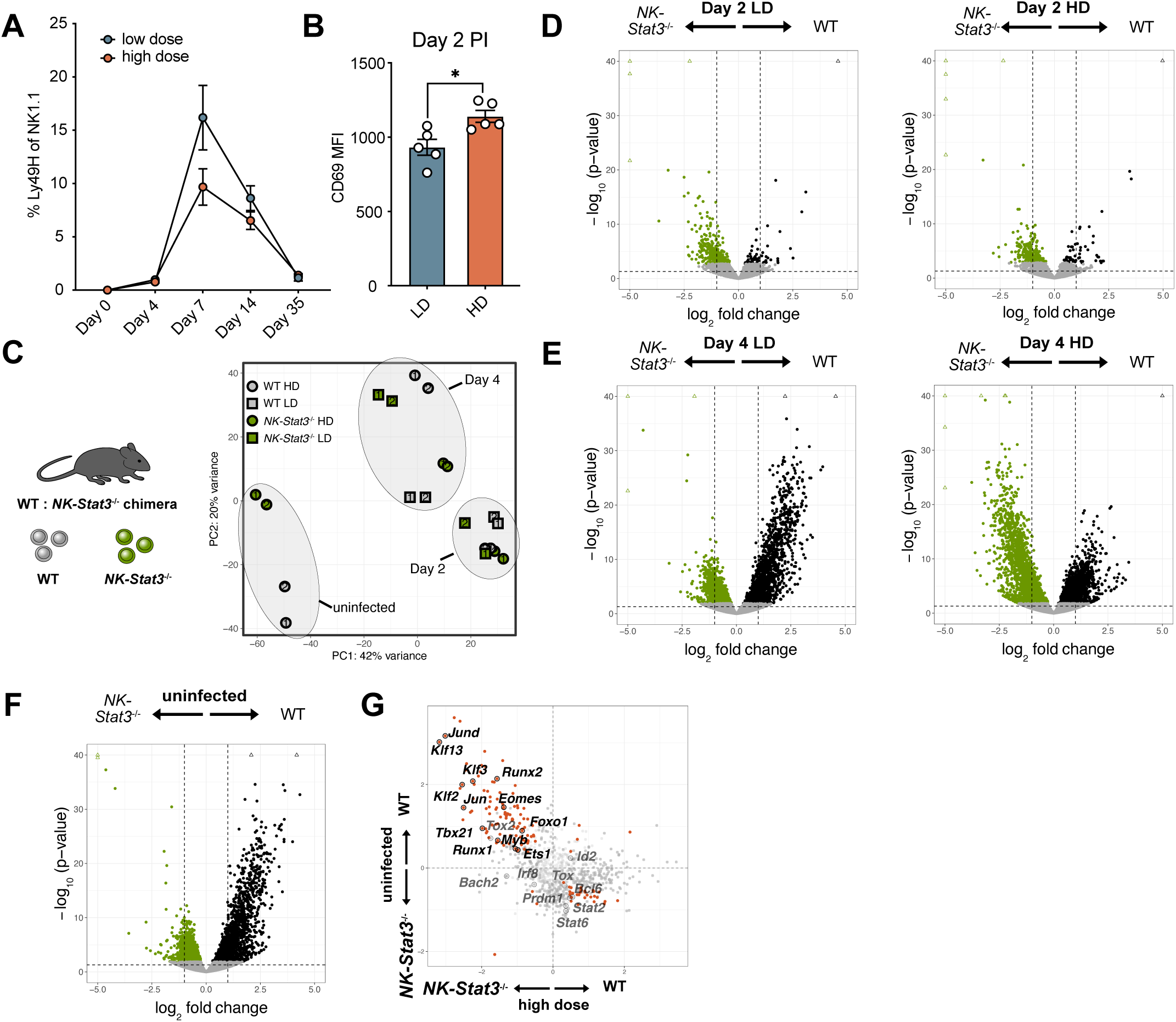
**(A)** Expansion kinetics of WT NK cells in low-dose and high-dose infection. **(B**) CD69 MFI of NK cells (not transferred) in low-dose and high-dose infection. **(C)** PCA plot of RNAseq experiment shown in Fig. 1D. **(D)** Volcano plot of RNA-seq from WT and *NK-Stat3^-/-^*NK cells on day 2 PI (low and high dose). **(E)** Volcano plot of RNA-seq from WT and *NK-Stat3^-/-^* NK cells on day 4 PI (low and high dose). NK cells were sorted from competitive WT:*NK-Stat3^-/-^* chimeric mice. Circles in color are significant DE genes. **(F)** Volcano plot of RNA-seq from WT and *NK-Stat3^-/-^*NK cells in uninfected mice. NK cells were sorted from competitive WT:*NK-Stat3^-/-^* chimera. Circles in color are significant DE genes. **(G)** Scatterplot of RNA-seq from high-dose MCMV infection (x-axis) and uninfected controls (y-axis). Circles in color are genes that are significantly DE at both contrasts. Highlighted are the transcription factor genes.

**Supp. Fig. 2:**
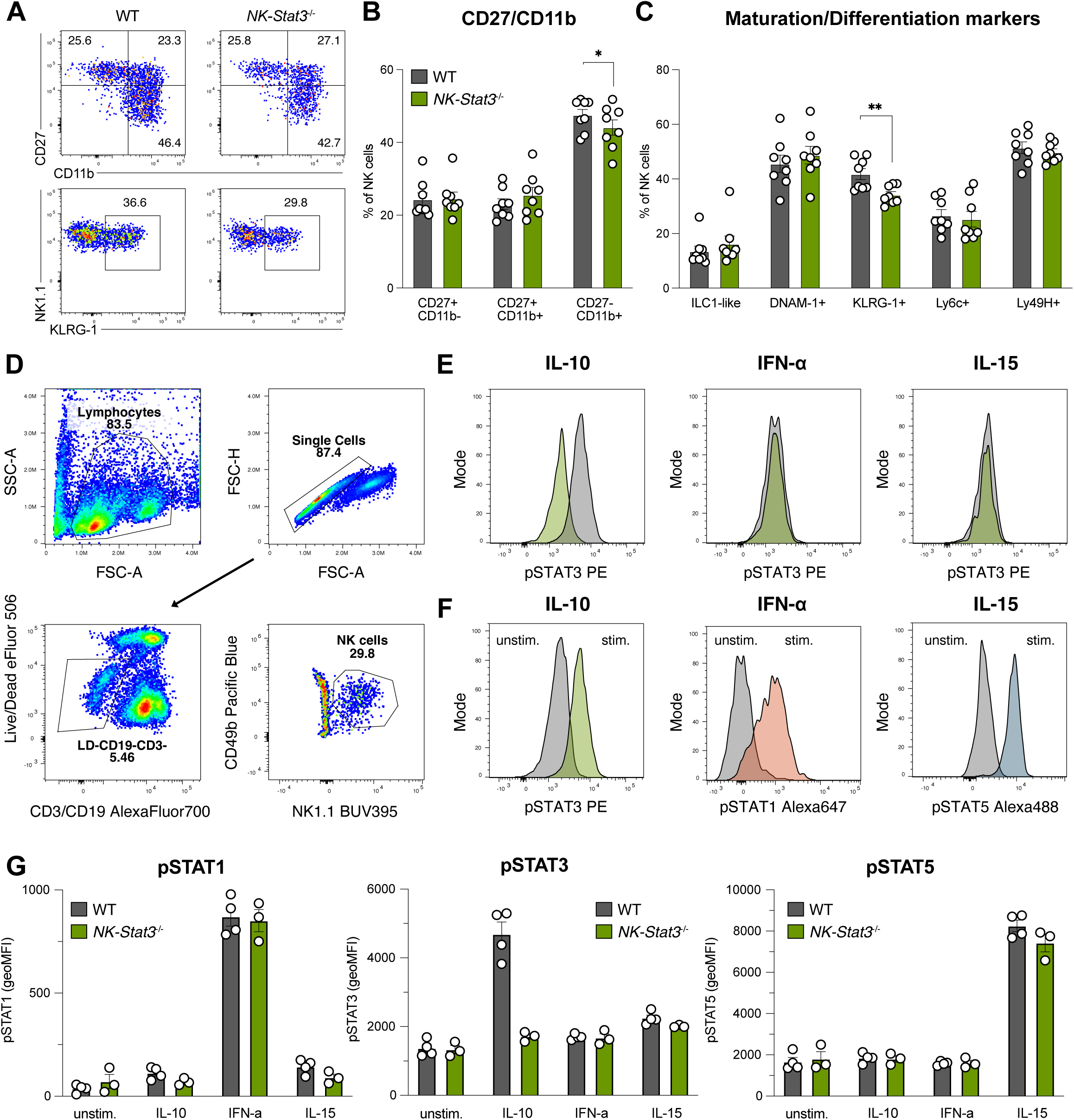
**(A)** Representative flow cytometry plots of maturation states of WT and *NK-Stat3^-/-^* NK cells under steady-state conditions cells were derived from competitive WT:*NK-Stat3^-/-^* chimera. **(B)** Quantification of maturation markers from (A) [top row]. **(C)** Quantification of maturation markers from A) [bottom row] and additional markers. **(D)** Representative gating for pSTAT staining in E). **(E)** pSTAT staining of WT and *NK-Stat3^-/-^*NK cells after stimulation with the indicated cytokines for 1h.

**Supp. Fig. 3:**
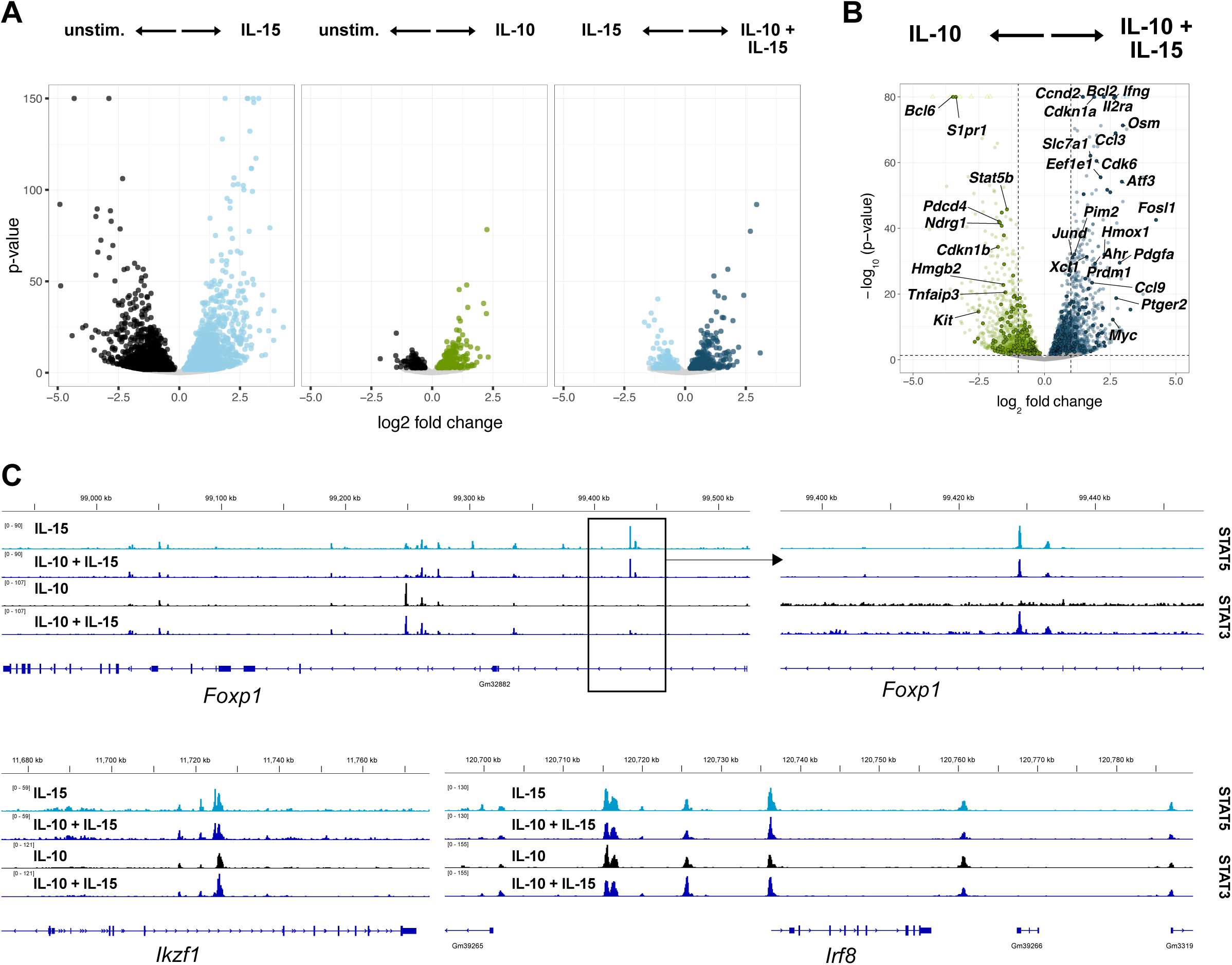
**(A)** Volcano plots for RNA-seq of NK cells stimulated with the indicated cytokines or unstimulated NK cells. Circles in color represent significant DE genes. **(B)** Volcano plot of NK cells stimulated with IL-10 + IL-15 versus IL-10. Circles in color are significant DE genes, circles in color with outlines are transcription factors, and gene name highlighted are from a cell proliferation pathway (see methods). **(C)** Representative tracks of STAT3 and STAT5 CUT&RUN in the indicated conditions. Boxed region in Foxp1 is zoomed in for clarity.

**Supp. Fig. 4:**
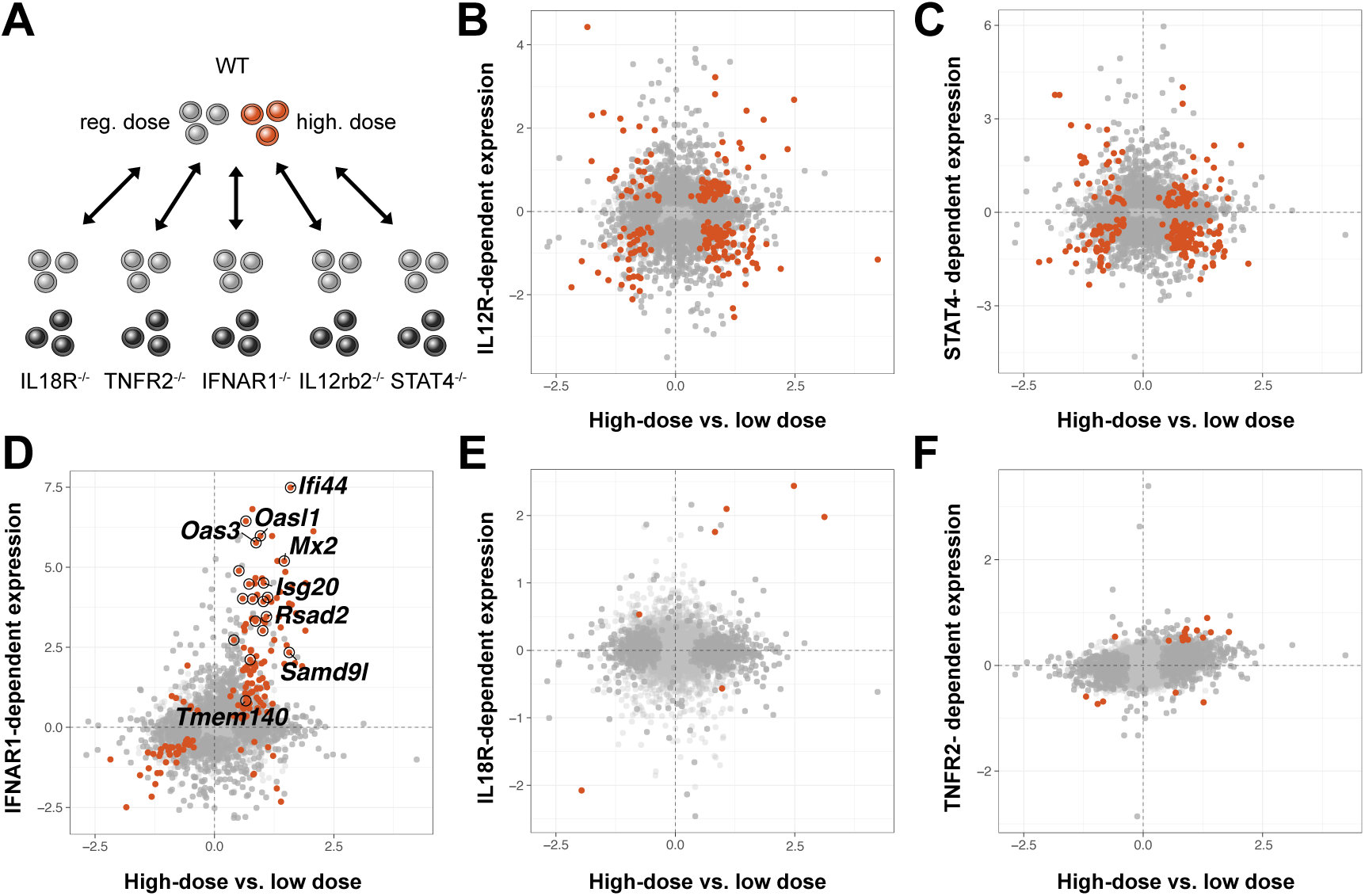
**(A)** Experimental schematic: Comparison of RNA-seq from day 2 PI from different cytokine receptor- and STAT-deficient to WT NK cells in HD versus LD infection. Comparison of WT NK cell in HD versus LD to **(B)** IL-12R-dependent genes, **(C)** STAT4-dependent genes, **(D)** IFNAR1-dependent genes, **(E)** IL18R-dependent genes and **(F)** TNFR2-dependent genes.

**Supp. Fig. 5:**
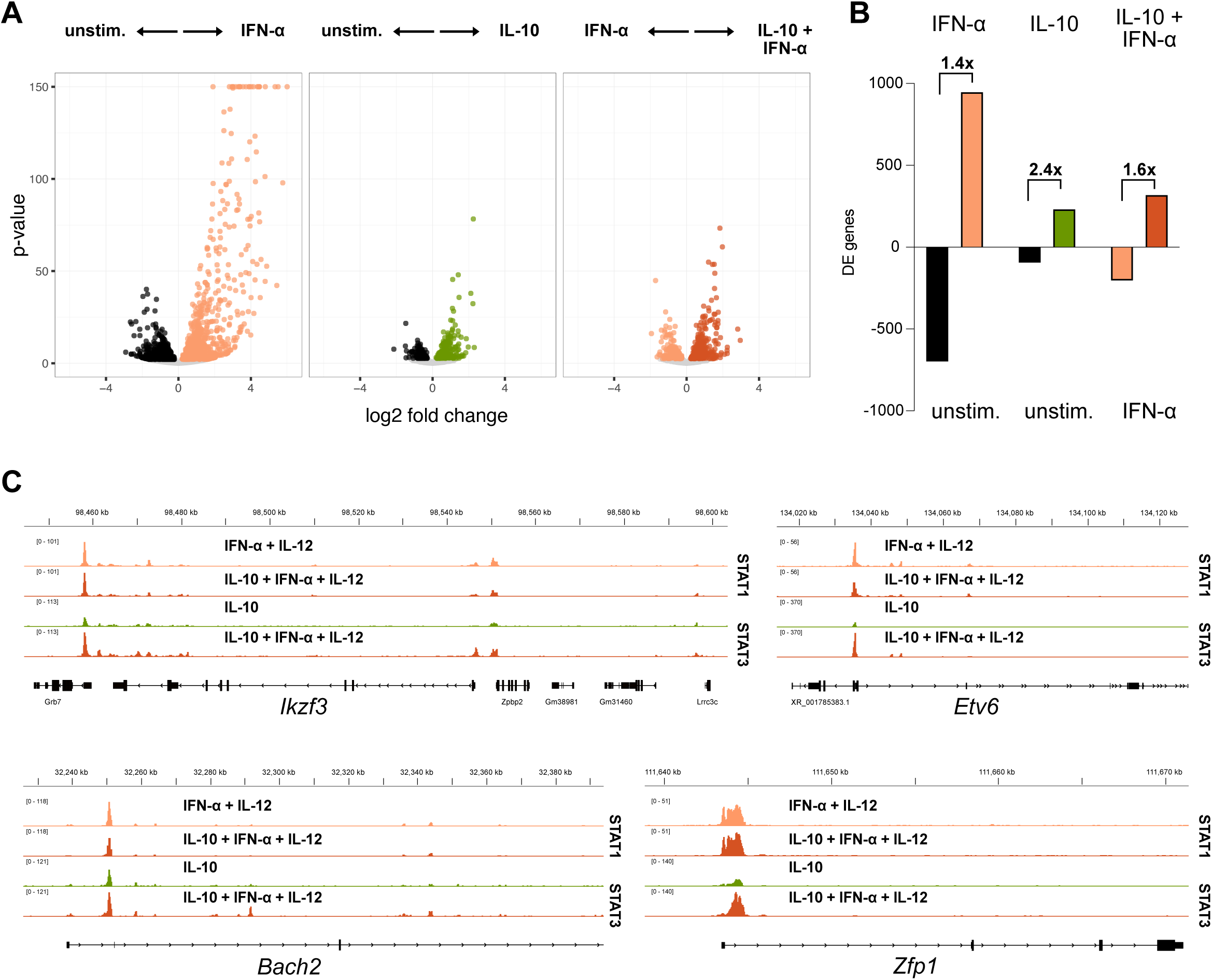
**(A)** Volcano plots for RNA-seq of NK cells stimulated with the indicated cytokines. Circles in color represent significant DE genes. **(B)** Quantification of significant up- or downregulated genes in conditions from (A). Values indicate (# upregulated DE genes) / (# downregulated DE genes). **(C)** Representative tracks of STAT1 and STAT3 CUT&RUN in the indicated conditions.

